# Morphological differences along the radial gradient of hippocampal area CA2 pyramidal neuron dendrites

**DOI:** 10.64898/2026.04.24.719171

**Authors:** Ivain Raslain, Ludivine Therreau, Vincent Robert, Houssam El Hariri, Vivien Chevaleyre, Peter Jedlicka, Hermann Cuntz, Rebecca A Piskorowski

## Abstract

Hippocampal area CA2 has recently emerged as a critical region for social recognition memory. Furthermore, this understudied region has been implicated in psychiatric diseases and neurodegenerative diseases. There has been accumulating evidence indicating that the pyramidal neurons (PNs) in area CA2 exhibit functional specializations that correlate with somatic position in stratum pyramidale (sp). In this study, we investigated the morphological differences in dendritic architecture of CA2 PNs with a focus on the radial gradient, i.e., along the deep–superficial axis of the sp. We conducted a comprehensive morphological analysis including Sholl intersection profiles, branching order distributions, root angle distributions, and dendritic cable lengths. We found that CA2 PNs have fewer oblique dendrites and a larger number of tuft-like dendrites as compared to CA1 PNs. Furthermore, within the CA2 population, we found that many of the dendritic structural features gradually changed along the radial axis from deep to superficial somatic location, indicating a continuum of dendritic morphology rather than two sharply defined subtypes of pyramidal neurons. This morphological characterization may serve as a starting point to better understand the corresponding functional organization of CA2. The gradual difference between deeper and superficial CA2 PNs suggests a continuum of their computational capabilities beyond two binary functional classes.

**In brief:** Using several methods, we examine the dendritic morphology of over 130 CA2 and CA1 pyramidal neurons and find that many properties such as the cable length and terminal numbers of the dendritic arbors vary as a with the location of the soma in the pyramidal layer.

**Highlights:**

- We use scholl analysis, graph theory and machine learning techniques to quantify the different dendritic morphologies of CA2 pyramidal neurons.
- Many properties of CA2 pyramidal neuron apical dendrites vary as a function of somatic location in the pyramidal layer.
- More superficial CA2 pyramidal neurons have longer oblique apical dendrites, and shorter tuft dendrites.

## Introduction

Memory formation and recall are essential for survival and forming relationships. Human and animal studies have established that the hippocampus is a fundamental structure for these functions (Squire and Wixted, 2011). The focus of this study is on hippocampal area CA2, a crucial region for social recognition memory (Hitti and Siegelbaum, 2014) found to be particularly vulnerable in numerous psychiatric disorders and neurodegenerative diseases (Chevaleyre and Piskorowski, 2016).

Recent studies that have recorded the activity of CA2 pyramidal neurons (PNs) during awake behavior have reported different activity patterns between deep (bordering *stratum oriens*, dCA2) and superficial (bordering *stratum radiatum*, sCA2) CA2 PNs. Specifically, in two studies using large scale high-density silicon probes to examine sharp-wave ripple (SWR) activity in all hippocampal subfields Oliva et al. (2016a,b), the burst index was higher in dCA2 as compared to sCA2. dCA2 PNs ramped up their action potential firing frequency preceding SWR activity in area CA1 and are then silent during the ripple (see also Kay et al., 2016). In contrast, sCA2 neurons fired in a phasic manner, with increased firing to the peak of the SWR followed by a strong inhibited response.

A novel CA2 intrahippocampal subcircuit important for memory consolidation was recently identified (Karaba et al., 2024). During non-rapid eye movement sleep, a subset of CA2 PNs fire long barrages of action potentials that were anti-correlated with SWRs in area CA1. CA1 neurons that increased action potentials during SWRs were inhibited during barrage firing. Interestingly, CA2 PNs that had the highest firing rate during barrage firing had somata located in the deep region of the pyramidal layer.

Further differences between deep and superficial CA2 PNs have been reported in *ex vivo* experiments examining cellular connectivity and physiology. Paired recordings of CA2 PNs revealed an increased probability of recurrent connectivity in superficial cells (Okamoto and Ikegaya, 2019). And lastly, studies using whole-cell recordings in acute brain slices showed an increased precedence of undergoing delta-opioid mediated inhibitory synaptic plasticity in superficial CA2 pyramidal neurons as compared to deep (Loisy et al., 2022).

The morphology of neurons is critical in shaping their electrophysiological properties (London and Häusser, 2005; Poirazi and Papoutsi, 2020). For example, the electrotonic distance and attenuation of passive propagation signals are influenced by dendritic length (Segev et al., 2003). And the action potential bursting in particular is thought to be crucially dependent on dendritic morphology (Duijnhouwer et al., 2001; van Elburg and van Ooyen, 2010; Cuntz et al., 2021). Thus, dendritic morphology should play a key role in determining the integrative properties of neurons in the hippocampus (Emri et al., 2001). Understanding these morphological and electrophysiological relationships in CA2 PNs is essential for elucidating the unique functional properties and information-processing mechanisms of the CA2 region within the hippocampus.

In this study, we reconstructed the complete apical and basal dendritic arbor of over 100 CA2 pyramidal neurons. We then employed the TREES toolbox (Cuntz et al., 2010) to analyze and differentiate the morphologies of CA2 PNs into two sub-populations: sCA2 and dCA2. We compared the morphologies of CA2 PNs with CA1 PNs to validate our methodology and then examined the dendritic morphology of CA2 PNs across somatic depth in the *stratum pyramidale*.

## Results

### Distinctions between CA1 and CA2 PNs

In order to check that CA2 PNs distinguish themselves from the very well-studied CA1 PNs in our hands, we first compared the two types of morphologies with our own methods using the exact same reconstruction pipeline for both cell types. Initially, we examined the Sholl intersection profiles (SIP; **Figure 1A–B**), to gain a comprehensive overview of the distinctions between the dendritic architecture of the different types of neurons (see **Methods** “Sholl analysis”). The SIP calculates the number of intersections between the dendritic tree and spheres of increasing radius centered on the neuron’s somata. This results in a one-dimensional representation of the morphology and was performed here separately for apical and basal dendrites. We conducted this analysis separately for the apical and basal segments, as discernible disparities between these segments were evident upon visual inspection of the neurons (**Figure 1A**). Apical dendrites exhibited bimodal distributions (**Figure 1B**, *Left*), with the first proximal peak corresponding to the *stratum radiatum* and the second distal peak to the *stratum lacunosum moleculare*. Notably, in CA1 PNs, the first proximal peak predominated over the second distal peak, whereas in CA2 PNs, this pattern was reversed. In the basal dendrites, the Sholl intersection profiles manifested as an unimodal curve with minimal distinctions between CA1 and CA2. SIP peaks in CA1 PNs occurred slightly earlier compared to CA2 PNs, accompanied by a shallower descending slope (**Figure 1B**, *Right*). These findings suggest a higher prevalence of apical oblique dendrites in CA1 PNs compared to CA2 PNs, while CA1 PNs exhibited fewer apical tuft dendrites relative to CA2 PNs.

**Figure 1.**
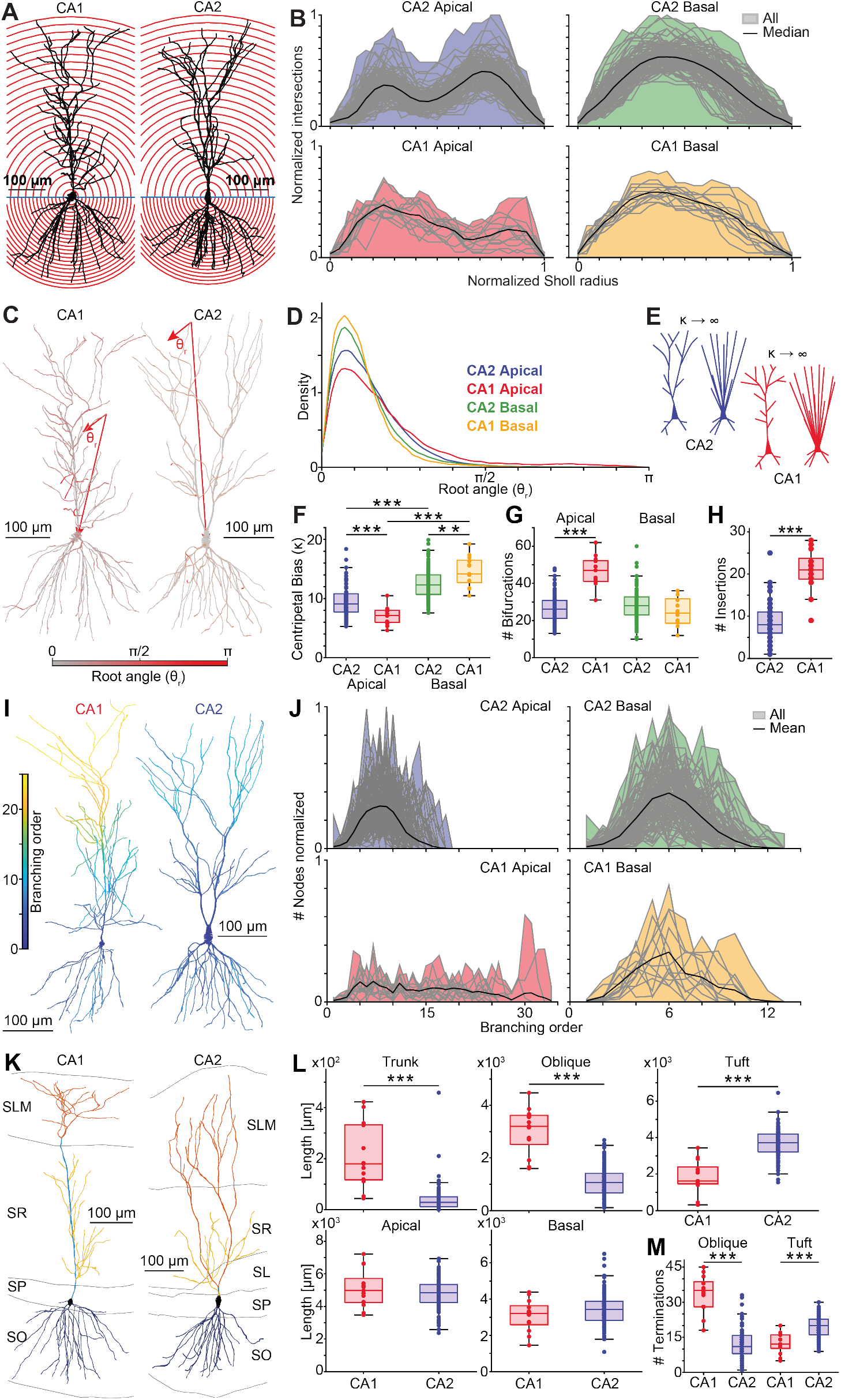
Morphological comparison of CA1 and CA2 PNs. **A**, Sholl analysis for sample reconstructed CA1 and CA2 PNs (see text for explanation). (See next page.) **Figure 1. (continued) B**, Overlaid Sholl intersections profile for apical and basal dendrites separately. Gray lines, individual trees; Black lines, median of average data; Shaded colored areas cover the maximal Sholl values for each radius value. **C**, Root angle for sample CA1 and CA2 PNs (see text). **D**, Distributions of root angles in CA1 and CA2 PNs for apical and basal dendrites separately. **E**, Sketch describing the impact of centripetal bias *κ* on morphology of PNs: As *κ* increases, root angle values tend to concentrate more tightly around 0. **F**, Estimated centripetal bias of the root angle distribution in the four conditions. **G**, Corresponding numbers of bifurcations. **H**, Number of obliques insertions. **I**, Branching orders in sample CA1 and CA2 PNs. **J**, Overlaid branch order distributions. **K**, Sample CA1 and CA2 PNs with color-coded sections and depiction of hippocampal layers. **L**, Cable length as it divides per section. **M**, Number of CA1 and CA2 termination points by section. Throughout the figure: Red – CA1 or CA1 apical depending on context; Blue – CA2 or CA2 basal depending on context; Green – CA2 basal; Yellow – CA1 basal.

We then calculated root angle distributions, that is, the distribution of angles corresponding to the angle that each dendritic segment deviates from a direct path to the soma (Bird and Cuntz, 2019). Flatter distributions indicate less optimized paths towards the soma while skewed distributions with higher centripetal bias *κ* indicate a more efficient signaling path towards the soma. When comparing the root angle distribution, we observed a significantly higher centripetal bias *κ*, for CA2 apical dendrites (mean *κ* = 9.44 ± 0.23) compared to CA1 apical dendrites (mean *κ* = 7.09 ± 0.43) (Wilcoxon U-test, p-value *<* 0.001) (**Figure 1C–F**). In contrast, for basal dendrites, CA2 (mean = 12.54 ± 0.24) exhibited a smaller *κ* value than CA1 (mean = 14.62 ± 0.70) (T-test, p-value *<* 0.001). Additionally, significant differences were found when comparing CA2 apical with basal dendrites (Wilcoxon U-test, p-value *<* 0.001), and CA1 apical with basal dendrites (T-test, p-value *<* 0.001). These observations indicate that the apical dendrites of CA1 PNs tend to spread more orthogonally from their main axis, represented by the trunk, compared to CA2 PNs.

We proceeded to examine the branching order (BO) distributions of both types of PNs (**Figure 1I–J**). Upon comparing the apical dendrites, it became evident that CA2 PN branching order distribution was significantly more concentrated around a smaller mean value, ranging between 7-9, with a maximum branching order of 18. Conversely, CA1 PN branching order distribution was more dispersed, lacking a clear mean, and with a maximum value of 34. This pattern aligns with the number of bifurcations observed in apical dendrites (**Figure 1G**), with CA2 PNs exhibiting a mean of 32.07 ± 0.71 bifurcations and CA1 PNs a mean of 46.77 ± 2.26 bifurcations (Wilcoxon U-test p-value *<* 0.001). Concerning the basal dendrites, the distributions appeared more similar, although CA1 PNs exhibited a smaller peak. The maximum branching order was 12, and a peak was observed at 6. However, the number of bifurcations is not significantly different between the two types of cell’s basal dendrites (CA2 mean = 28.12 ± 0.73, CA1 mean = 24.77 ± 2.16, Wilcoxon U-test p-value = 0.092).

Additionally, we looked at the number of insertions of oblique dendrites along the main apical dendrites (**Figure 1H**), finding that CA2 PNs displayed fewer oblique insertions than CA1 (CA2 mean = 8.37 ± 0.37, CA1 mean = 20.69 ± 1.45, Wilcoxon U-test p-value *<* 0.001).

We then analyzed the cable length of distinct sections of PNs (**Figure 1K–L**), including the apical, basal, trunk, obliques, and Tuft/Tuft-like sections (see **Methods** “Defining pyramidal cell compartments”). CA1 PNs exhibited a greater trunk cable length (mean = 0.210 ± 0.035 mm) compared to CA2 (mean = 0.039 ± 0.005 mm) (Wilcoxon U-test p-value *<* 0.001) and a higher oblique cable length (CA1 mean = 3.037 ± 0.244 mm, CA2 mean = 1.074 ± 0.054 mm, Wilcoxon U-test p-value *<* 0.001) (**Figure 1L**). Conversely, for the distal tuft dendrites, CA2 PNs had a greater cable length (mean = 3.656 ± 0.071 mm) compared to CA1 PNs (mean = 1.842 ± 0.250 mm) (Wilcoxon U-test, p-value *<* 0.001). However, no significant differences were observed for the apical part (CA1 mean = 5.093 ± 0.322 mm, CA2 mean = 4.772 ± 0.084 mm, T-test p-value = 0.32). Likewise, in the basal dendrites, no significant difference in cable length was noted, with CA1 displaying a lower cable length (mean = 3.085 ± 0.241 mm) than CA2 (mean = 3.446 ± 0.085 mm) (T-test, p-value = 0.178). Furthermore, **Figure 1M** illustrates a significant difference in the number of dendrite terminations, reflecting the varying number of dendrites per layer/section of the neurons, between CA1 oblique (mean =33.54 ± 2.18) and CA2 oblique (mean = 11.94 ± 0.60) (Wilcoxon U-test, p-value *<* 0.001), and between CA1 tuft (mean = 12.69 ± 1.23) and CA2 tuft-like (mean = 19.44 ± 0.41) (T-test, p-value *<* 0.001).

### Comparison between deep and superficial CA2 pyramidal neurons (PNs)

To test our hypothesis of the existence of two morphological subtypes as in CA1, we replicated our analysis to explore morphological disparities within CA2 PNs, classifying them as either deep (dCA2) or superficial (sCA2) based on the location of their soma in *stratum pyramidale* (**Figure 2A**, see **Methods** “Classifying neurons”).

**Figure 2.**
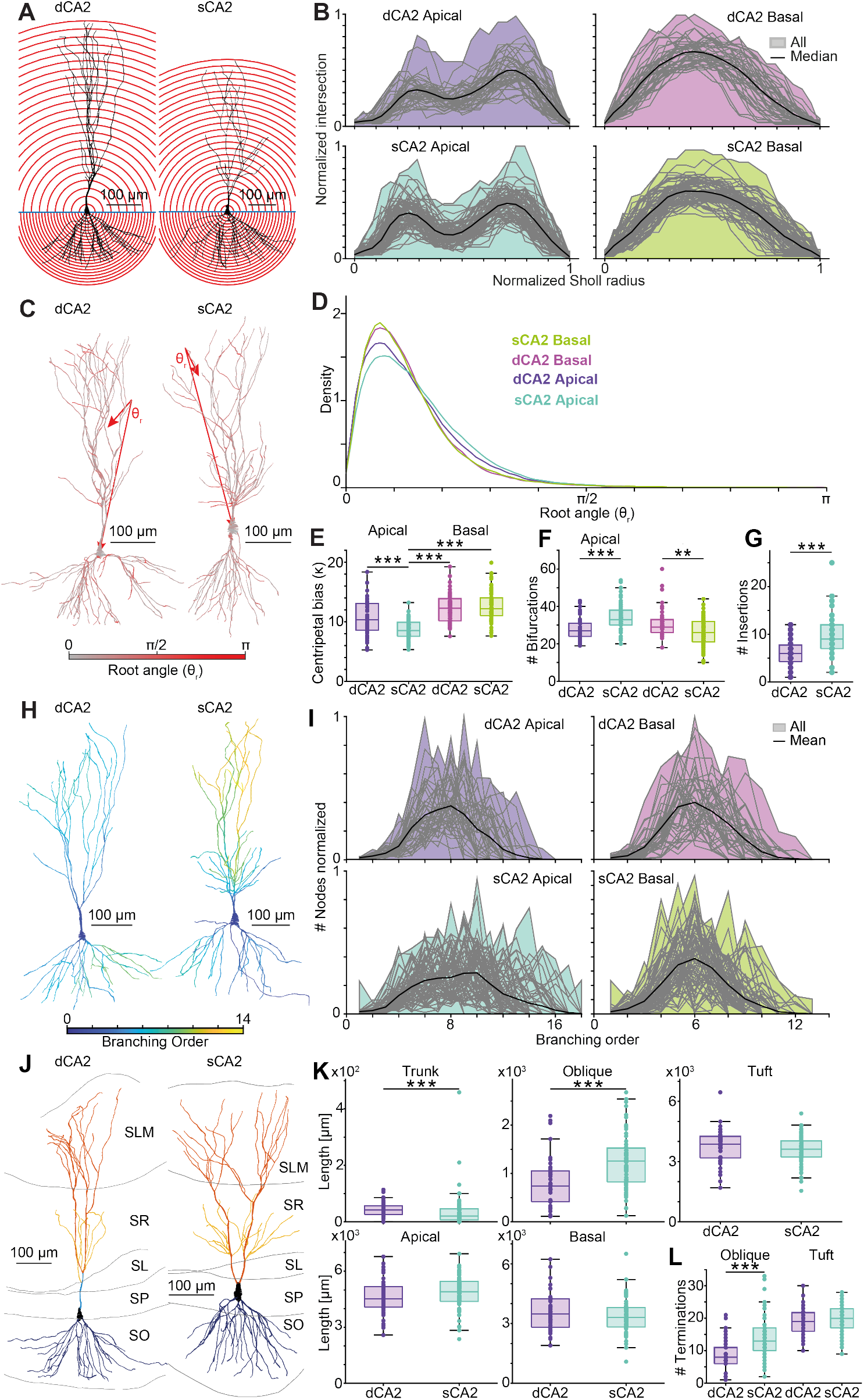
Comparative analysis of deep CA2 (dCA2) and superficial CA2 (sCA2) PNs. Similar layout is in **Figure 1** but comparing dCA2 and sCA2 PNs. **A**, Sholl analysis for sample dCA2 and sCA2 PNs. **B**, Overlaid Sholl intersection profiles. **C**, Root angles for sample dCA2 and sCA2 PNs. **D**, Distribution of root angles for all dCA2 and sCA2 PNs separated into their basal and apical dendrites. **E**, Estimated centripetal bias *κ*. **F**, Number of bifurcations. **G**, Number of obliques insertions. **H**, Branching orders for sample dCA2 and sCA2 PNs. **I**, Overlaid branching order distributions for all dCA2 and sCA2 PNs separated into their basal and apical dendrites. **J**, sample dCA2 and sCA2 PNs with color-coded sections and depiction of hippocampal layers. **K**, Cable length as it divides per section. **L**, Number of dCA2 and sCA2 termination points by section. Throughout the figure: Purple – dCA2 PNs or their apical dendrites depending on context; Cyan – sCA2 PNs or their apical dendrites depending on context; Magenta – dCA2 PN basal dendrites; Light green – sCA2 PN basal dendrites.

Basal sCA2 PN dendrites appeared to exhibit a slightly higher peak in the normalized Sholl intersection profile (**Figure 2B**). However, when examining the apical dendrites, the second peak corresponding to distal dentrites appeared at the same normalized Sholl radius with similar height and width, while differences were observed in the first peak corresponding to proximal dendrites. Indeed, sCA2 PNs exhibited a higher and earlier proximal dendrite peak. These results suggest that sCA2 PNs have a more proximal main bifurcation point and more oblique dendrites, whereas dCA2 PNs have a higher basal dendritic density.

We observed that apical dCA2 PNs exhibited a more concentrated distribution of root angles compared to apical sCA2 (dCA2 mean *κ* = 10.67 ± 0.45, sCA2 mean *κ* = 8.73 ± 0.20, Wilcoxon U-test p-value *<* 0.001) (**Figure 2D-E**). However, no significant differences were observed in the root angles of basal dendrites (dCA2 mean *κ* = 12.30 ± 0.40, sCA2 mean *κ* = 12.69 ± 0.30, T-test p-value = 0.86). Nevertheless, within the superficial PNs, significant differences were detected in root angles of different sections (T-test, p-value *<* 0.001). These findings indicate that while basal dendrites adopt a similar pattern of occupying *stratum oriens* in both dCA2 and sCA2 PNs, apical dendrites of PNs exhibit greater spreading in sCA2 than in dCA2.

Upon examining the branching order distribution, we report a noticeable difference between dCA2 and sCA2 PN apical but not basal dendrites (**Figure 2I**). For the apical dendrites, the branching order distribution appeared more concentrated in dCA2 PNs, reaching a maximum value of 14, compared to superficial ones with a maximum value of 17. Furthermore, there were fewer bifurcations in dCA2 PNs (mean = 28.12 ± 0.92) compared to sCA2 (mean = 34.43 ± 0.89) (Wilcoxon U-test, p-value *<* 0.05). Conversely, in basal dendrites, dCA2 PNs exhibited a greater number of bifurcations (mean = 31.19 ± 1.20) than sCA2 PNs (mean = 26.29 ± 0.85) (Wilcoxon U-test, p-value *<* 0.05)(**Figure 2F**). These results indicate that sCA2 PN apical dendrites possess more oblique dendrites than dCA2 PNs. The difference in basal dendrites seems to be in the more distal branches, as evidenced by the lack of notable impact on the branching order distribution. Additionally, dCA2 PNs (mean = 6.21 ± 0.46) display less oblique insertion than sCA2 PNs (mean = 9.65 ± 0.48) (Wilcoxon U-test, p-value *<* 0.001) (**Figure 2G**).

In comparing the cable lengths for each dendritic compartment (**Figure 2J–K**), we did not find significant differences in the tuft (dCA2 mean = 3.772 ± 0.129 mm, sCA2 mean = 3.587 ± 0.082 mm, T-test p-value = 0.231), apical (dCA2 mean = 4.607±0.145 mm, sCA2 mean = 4.871±0.103 mm, Wilcoxon U-test p-value = 0.142), and basal (dCA2 mean = 3.574 ± 0.149 mm, sCA2 mean = 3.369 ± 0.102 mm, T-test p-value = 0.261) dendrites. However, in the trunk we found a higher cable length for dCA2 compare to sCA2 (deep mean = 0.044 ± 0.004 mm, sCA2 mean = 0.035 ± 0.007 mm, Wilcoxon U-test p-value *<* 0.001), and in the oblique dendrites, we found that sCA2 PNs had a higher cable length (mean = 1.245 ± 0.064 mm) than dCA2 PNs (mean = 0.788 ± 0.081 mm) (T-test, p-value *<* 0.001). Additionally, the number of terminations, as shown in **Figure 2K**, was higher for oblique sCA2 PNs (mean = 14.08±0.74) than for dCA2 PNs (mean = 8.35 ± 0.75) (Wilcoxon U-test, p-value *<* 0.005), but not for the tuft dendrites where they were not significantly different (dCA2 mean = 18.84 ± 0.68, sCA2 mean = 19.79 ± 0.51, T-test p-value = 0.231). These results indicate that sCA2 PNs have a greater cable length in the oblique section due to more numerous dendrites.

### Non-biased classification of CA2 PN subpopulations

To enhance the robustness of our morphological classification, we used machine learning classifiers and clustering algorithms. To accomplish this, we computed various morphological measures previously illustrated in **Figure 1** and **2** (see Methods “Morphological classification using machine learning”) and attempted to replicate the methodology outlined by (Vasques et al., 2016). Initially, we fine-tuned algorithm parameters by classifying CA1 and CA2 neurons. The results demonstrated satisfactory accuracy scores across all algorithms (**Figure A**, *Red line*). This indicated that our morphological measurements enabled confident classification of CA1 and CA2 PNs. However, extending this classification to distinguish between dCA2 and sCA2 PNs yielded less favorable results. Accuracy scores hovered around 0.7, indicating moderate predictive power but suggesting room for improvement.

Additionally, we used unsupervised clustering algorithms and compared their accuracy using the V-Measure an entropy-based measure, harmonically combining homogeneity and completeness scores (**Figure B** Rosenberg and Hirschberg, 2007). For all algorithms used, clustering CA1 and CA2 neurons gave high V-measure, with a slight decrease for affinity propagation algorithms (red line). Again, applying the same fine-tuned clustering algorithms to dCA2 versus sCA2 PNs was not satisfactory. Indeed, with no regard to the number of clusters asked, the V-measures were between 0 and 0.2, indicating a low validity of these clustering solutions. In addition, while CA1 and CA2 PN morphologies separated in principal components space, dCA2 and sCA2 morphological differences seemed more gradual with soma depth (**Figure C**).

While the previous analysis in figures 1 and 2 is relevant, because they represent both ends of the distribution of somatic locations. These unbiased results led us to hypothesize a continuous variation along the radial hippocampal axis rather than a strict binary classification into superficial and deep subpopulations in CA2.

### Correlation of soma location with dendritic architecture

To evaluate our new hypothesis, we performed a correlation analysis between various morphological characteristics and the radial location of the PN soma (**Figure 4A-D**), focusing on each defined region of the neuron (see **Methods** “Defining pyramidal cell compartments”).

**Figure 3.**
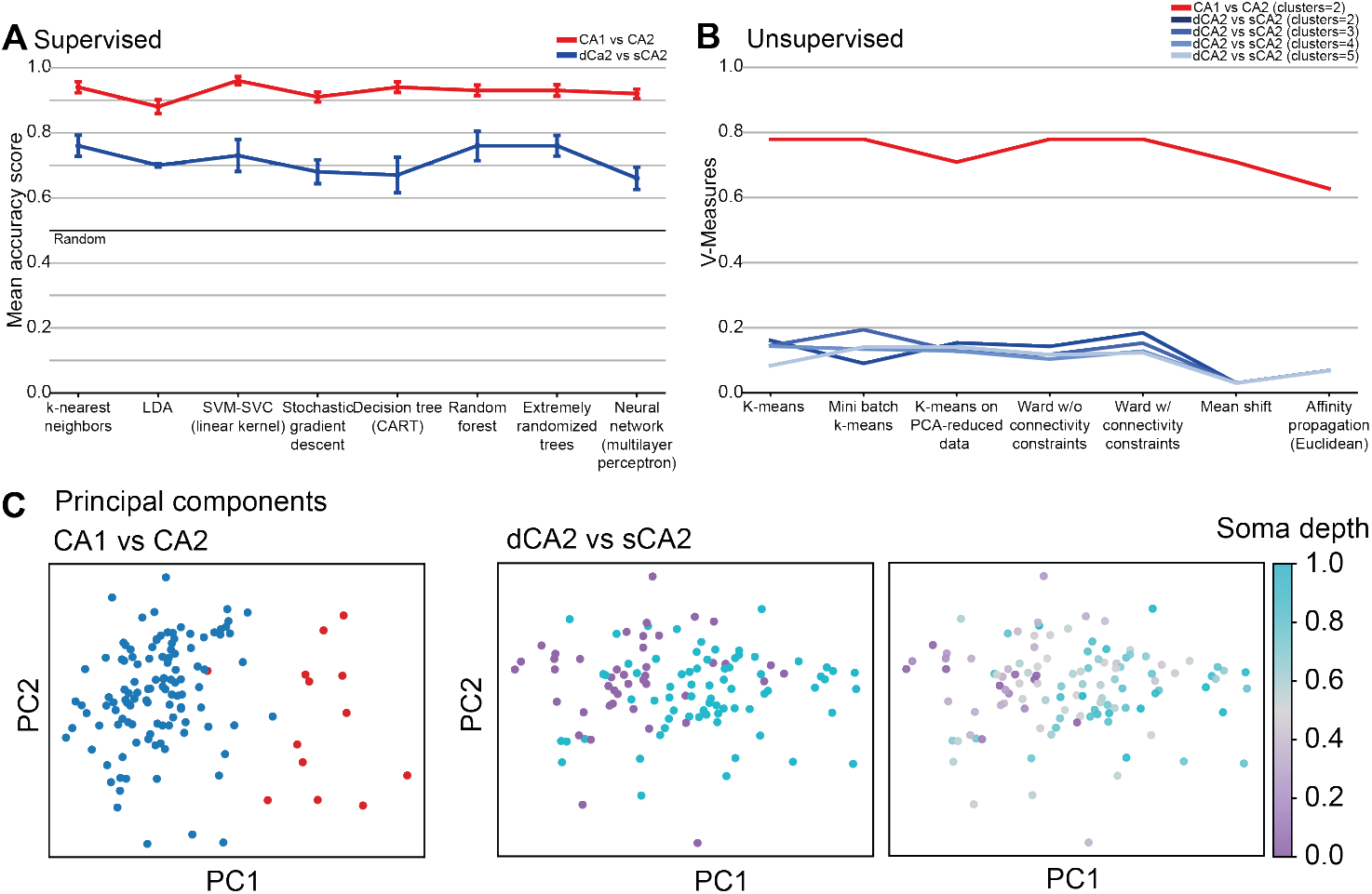
Morphological classification via supervised and unsupervised algorithms. **A**, Mean accuracy scores for supervised classification algorithms: k-nearest neighbors, linear discriminant analysis, support vector machine with a linear kernel, stochastic gradient descent, classification and regression trees, random forest, extremely randomized trees, and multilayer perceptron. The red line indicates scores for classifying CA1 and CA2 PNs, while the blue line represents scores for distinguishing between dCA2 and sCA2 PNs. Error bars represent the standard error of the mean (SEM). **B**, V-measures for unsupervised clustering algorithms: k-means, mini batch k-means, PCA-reduced k-means, Ward’s method (without connectivity constraints), Ward’s method (with connectivity constraints), mean shift, and affinity propagation (Euclidean). The red line depicts scores for discriminating CA1 and CA2 PNs into two clusters, while the blue lines represent scores for discriminating dCA2 and sCA2 PNs into two, three, four, and five clusters. Mean shift and affinity propagation algorithms did not allow for the specification of a desired number of clusters. **C**, The morphometrics for CA1 vs CA2 (*left*) and dCA2 vs sCA2 (*right*) plotted using principal component analysis (PCA) including a version with colored soma depth (*far right*).

**Figure 4.**
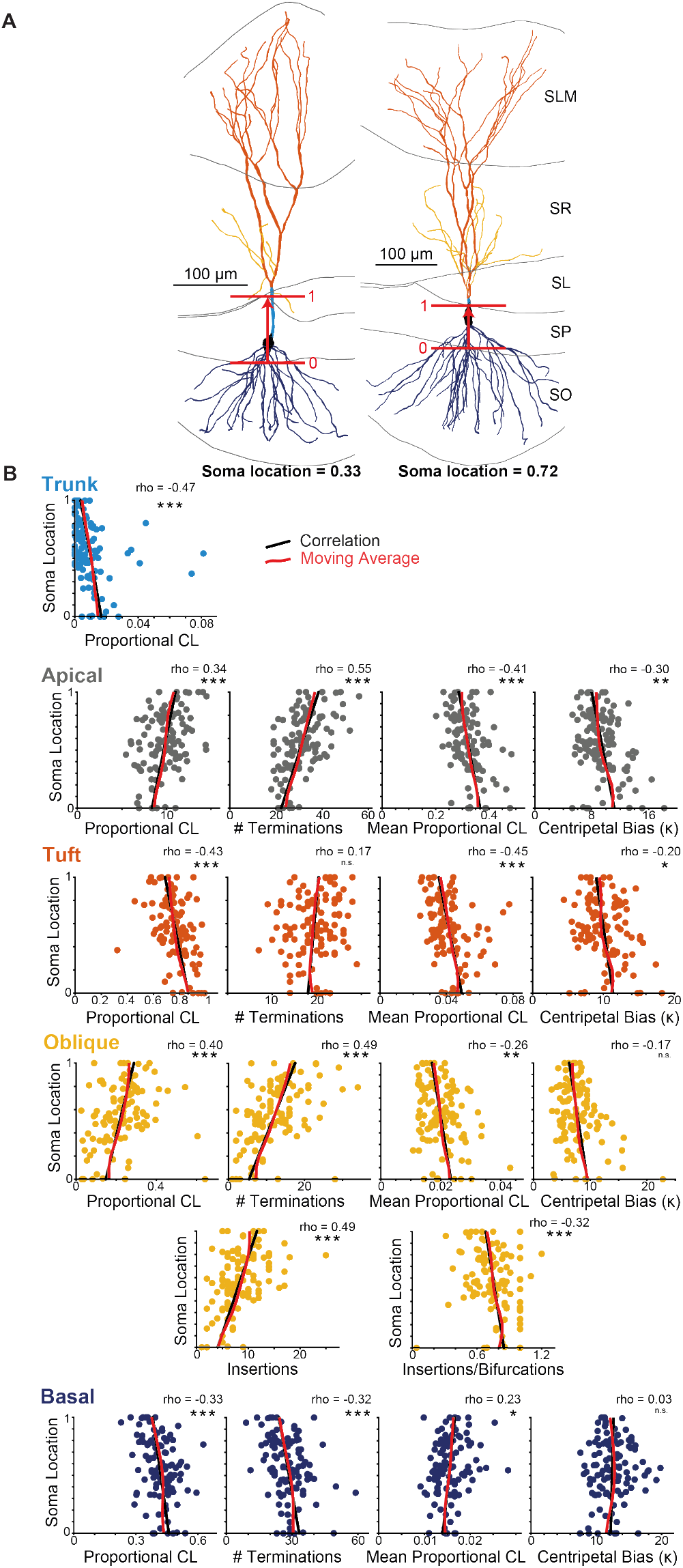
Depth-dependency of morphological measures in CA2 PNs. **A**, Two sample PNs with color-coded dendritic morphology classifications: Trunk (light blue), apical (gray), basal (dark blue), tuft (orange), oblique (yellow). Hippocampal lamina are diagramed, SO, stratum oriends, SP, stratum pyramidale, SL stratum lucidum, SLM, stratum lacunosum moleculare. Red lines illustrate the limits of soma radial location, with the value given for the two examples. **B**, Soma location as a function of morphological measures for the different dedritic regions. The morphological measaures are: proportional cable length (CL), number of terminations, mean proportional cabel length, centripetal bias, number of oblique insertions and oblique insertions ratio. Straight black lines represent correlation, while the red line illustrates the moving average. Correlation coefficients are displayed. (∗) indicates *p <* 0.05, (∗∗) indicate *p <* 0.01, and (∗ ∗ ∗) indicate *p <* 0.001.

As expected, a negative correlation was found between radial soma location and trunk PCL (*ρ* = −0.47, p-value *<* 0.001), indicating a shorter trunk as the soma is located more superficially. Because the trunk dendrite needs to traverse the pyramidal layer before branching in *stratum radiatum*, this negative correlation is a natural consequence of the somatic location and thickness of the pyramidal layer.

For apical dendrites, we found that radial soma location was positively correlated with the proportional cable length (PCL) (*ρ* = 0.34, p-value *<* 0.001) and the number of terminations (*ρ* = 0.55, p-value *<* 0.001), while it was negatively correlated with the mean PCL (*ρ* = −0.41, p-value *<* 0.001). Additionally, a significant negative correlation was observed with centripetal bias (*ρ* = −0.30, p-value *<* 0.01). These results suggest that neurons with more superficial soma develop more extensive and intricate dendritic arbors with a more outward-directed growth pattern.

For the tuft branches, radial soma location was negatively correlated with PCL (*ρ* = −0.43, p-value *<* 0.001), mean PCL (*ρ* = −0.45, p-value *<* 0.001), and centripetal bias (*ρ* = −0.20, p-value *<* 0.05). Despite this, there was no significant correlation with the number of terminations (*ρ* = 0.17, p-value = 0.07), indicating a reduction in proportional length and a more dispersed spatial distribution as the soma is located more superficially.

In the case of oblique dendrites, radial soma location showed a positive correlation with PCL (*ρ* = 0.40, p-value *<* 0.001), the number of terminations (*ρ* = 0.49, p-value *<* 0.001), and the number of insertion (*ρ* = 0.49, p-value *<* 0.001). In parallel, it showed a negative correlation with the mean PCL (*ρ* = −0.26, p-value *<* 0.01), and the ratio of insertions to the total number of bifurcations (*ρ* = −0.32, p-value *<* 0.001). In contrast, no significant correlation was found between radial soma position and centripetal bias (*ρ* = −0.17, p-value = 0.07). These findings suggest that more superficially located soma are associated with an increased number of oblique dendrites extending into *stratum radiatum*. This increase appears to be twofold: not only is there a higher number of dendrites branching off from the apical trunk and tuft compartments, but the oblique dendrites themselves also exhibit a greater degree of arborization. The negative correlation of ratio of insertions to the total number of bifurcations indicates that the increase in branching within the oblique dendrites may contribute more significantly to the overall complexity than the increase in the number of insertions alone.

Regarding the basal dendrites, no significant correlation was found between soma location and centripetal bias (*ρ* = 0.04, p-value = 0.81). However, a negative correlation with PCL (*ρ* = −0.33, p-value *<* 0.001) and with the number of terminations (*ρ* = −0.32, p-value *<* 0.001) and a positive correlation with mean PCL (*ρ* = 0.23, p-value *<* 0.05) has been found, implying that CA2 PNs with more superficial soma locations have fewer but longer basal dendritic branches.

## Discussion

In this study, we have extensively and systematically described the dendritic morphologies of reconstructed mouse hippocampal CA2 PNs based on various features, including Sholl intersection profile, root angle distribution, centripetal bias, number of bifurcations, number of oblique insertions, branching order distribution, and partial cable length. Using methodological segmentation of neuronal compartments, we identified and quantified morphological distinctions between CA1 and CA2 PNs. In CA1 PNs, dendritic branching typically follows a long apical trunk extending to the *stratum lacunosum moleculare*, where it branches into numerous tuft dendrites. In contrast, CA2 PNs exhibit a much shorter apical trunk that terminates in the *stratum lucidum* or early in the *stratum radiatum*, where it branches into tuft-like dendrites. These distal dendrites —sometimes referred to as secondary dendrites or tuft dendrites (Ishizuka et al., 1995)— are more numerous and longer in CA2 PNs than in CA1 PNs.

Along the apical trunk of CA1 PNs, we observed a large number of oblique dendrites oriented roughly orthogonal to the trunk. This dense distribution of oblique dendrites likely contributes to the lower centripetal bias that we observed in CA1 apical dendritic arbors. In contrast, CA2 PNs possess fewer oblique dendrites. Those that are present are oriented toward the *stratum lacunosum-moleculare* or run parallel to the main trunk, resulting in a larger centripetal bias. From these measures, seems that CA2 PNs prioritize conduction by minimizing total dendritic length, as individual dendrites are taking a shorter path to the soma (Bird and Cuntz, 2019).

We also identified differences in the number of oblique dendritic insertions between CA1 and CA2, a measure distinct from either the total number or total length of oblique dendrites. While dendrite number and length inform about potential synapse distribution and density, the number of insertions provides insight into the capacity for shunting distal synaptic inputs as they propagate toward the soma. Our results are in agreement with the experimental and computational studies demonstrating a 5- to 6-fold larger somatic EPSP from distal synaptic input as compared to inputs in the same dendritic compartment in CA1 PNs (Sun et al., 2014; Srinivas et al., 2017; Chevaleyre and Siegelbaum, 2010). This difference arises from dendritic morphology, a more prominent effect of dendritic Ih in CA1 than in CA2, and a threefold higher number of EC synaptic contacts onto the distal dendrites of CA2 PNs.

We found that the basal dendrites of CA2 PNs are different from CA1 in both length and the radial angle distribution. The physiological consequences of these differences have yet to be explored. In CA1, the synaptic inputs at these dendrites are primarily coming from CA2 PNs, and to a smaller fraction, CA3 and the septum. In area CA2, there is much less known about the inputs arriving at this compartment, but it is likely the CA2 recurrent connections, CA3 and septal inputs.

### Beyond binary classes

We then applied similar methods to compare deep CA2 (dCA2) and superficial CA2 (sCA2) PNs sub-populations of CA2, which we have formerly classified by their soma location in the *stratum pyramidal*. Indeed, by investigating the Sholl intersection profiles, we observed a higher proximal mode corresponding roughly to the *stratum radiatum* for sCA2 apical dendrites. Confirmed by higher maximum branching order, more numerous bifurcations, larger oblique cable length, number of oblique terminations and a difference in the root angle distributions. Furthermore, we observed a decrease in tuft-like centripetal bias in CA2 PNs from deep to superficial somatic location, suggesting a potential increase in conduction delay but a reduction in the metabolic cost associated with dendritic growth Bird and Cuntz (2019). This could allow broader coincidence detection for inputs arriving at different times. Additionally, we found an increase in the number of oblique dendrites, which could enhance their shunting effect on distal synaptic integration, as suggested by Piskorowski and Chevaleyre (2012), or alternatively, indicate a specialization for processing more proximal inputs, such as ipsilateral CA3 Schaffer collateral projections resulting in a stronger inhibition (Piskorowski and Chevaleyre, 2012; Chevaleyre and Siegelbaum, 2010).

Consistently, there is evidence from *in vivo* and *ex vivo* electrophysiology measurements that deep and superficial CA2 PNs have different firing properties and are potentially recruited at different times in hippocampal memory processes. Specifically, measurements performed with high density arrays of silicon probes have revealed two functional populations of CA2 PNs. There are “ramping cells” that are located in deeper pyramidal layers and increase their firing prior to ripple activity in area CA1, and then rapidly become silent during the ripple (Oliva et al., 2016a,b) . It is likely these same deep “ramping” CA2 cells that show peculiarly firing activity during spatial navigation, with increased firing during immobility (Kay et al., 2016). Furthermore, during sleep, deep CA2 ramping cells undergo prolonged “barrage” firing activity during non-rapid eye movement sleep (Karaba et al., 2024). In contrast, superficial CA2 PNs, or “phasic” cells undergo an increase in firing that proceeds and continues throughout ripple events in CA3 and CA1 (Oliva et al., 2016a,b). Hippocampus, 26, 1593–1607.) Recordings from pairs of CA2 pyramidal cells indicate that superficial cells may have a higher probability of being recurrently connected to other CA2 cells (Okamoto and Ikegaya, 2019). Lastly, we have shown that superficial CA2 pyramidal cells are much more likely to undergo endocannabinoid-mediated synaptic plasticity at inhibitory inputs in acute slice preparations that is critical for social recognition memory (Loisy et al., 2022).

Our results indicate that there are not two separable populations of CA2 PNs. Rather, we observe a gradient of differences along the radial axis. Gradients of dendritic structure have been described in area CA1 along the major physical axes of the hippocampus. For example, lateral CA1 PNs located closer to CA3 contain more basal dendrites and fewer apical tuft dendrites, while those located closer to the subiculum (i.e., more medially) contain fewer basal dendrites and more apical tuft dendrites (Graves et al., 2012). Similarly, dCA1 PNs have larger basal dendritic trees than sCA1 PNs (Lee et al., 2014). Likewise, pyramidal neurons in the dorsal hippocampus have more branches than their counterparts in the ventral hippocampus (Dougherty et al., 2012).

In CA1, 265 genes exhibited greater than twofold differential expression between CA1 pyramidal neurons in dorsal and ventral hippocampus (Cembrowski et al., 2016a). Consistent with our observations, the differences in gene expression in CA1 occur along gradients rather than as discrete cell populations. A separate study involving single-cell RNA sequencing also revealed gradients of gene expression (Yao et al., 2021; Zeisel et al., 2015, 2018). Gradients of gene expression, morphological and physiological properties generalize to other hippocampal areas, including CA3 and the dentate gyrus (Cembrowski et al., 2016b).

### Comparison with previous work

We did not find four possible subtypes as presented by Helton et al. (2019). One explanation would be that our dendritic reconstructions were performed from cells recorded from relatively thick transverse hippocampal slices. In coronal brain slice preparations, CA2 PN dendrites are unquestionably cut during the slicing process to a much greater extent, and will result in incomplete dendritic morphologies. Our results are consistent with those of Srinivas et al. (2017), who also based their analysis on mouse dendritic reconstructions prepared in a similar way.

## Conclusion

Building on the findings from this study, future work should focus on elucidating the functional implications of the morphological gradient observed within the CA2 PN population. Specifically, it would be beneficial to investigate how these morphological variations correlate with differences in electrophysiological properties and synaptic connectivity. Additionally, expanding the analysis to include a larger sample size and more refined classification techniques could help in better distinguishing the subpopulations within area CA2. Understanding these nuances could provide deeper insights into the role of CA2 in hippocampal circuits and its contribution to cognitive functions such as memory and social behavior.

## Acknowledgments

We are grateful to Tristan Stöber for comments on the manuscript. This work was supported by Fondation pour la Recherche Médicale EQU202003010457 to RAP and French National Research Agency ANR-18-CE370020-01, ANR-18-CE16-0006, ANR-21-CE16-0021-03 and ANR-23-CE14-0004 to RAP and by the Federal General Institute for Risk Assessment, Grant Agreement Number 60-0102-01.P636 to HC and PJ. IR and VR were recipients of PhD fellowships from the French Ministry of Research and Higher Education. We would like to acknowledge Julie Cognet for technical assistance. The authors declare to have no competing financial interests.

## Author contributions

VR, LT, HE, VC and RAP performed whole cell recordings, histology, confocal imaging and Neurolucida reconstructions. RAP, PJ, HC and IR designed the study. IR performed the simulations and analysed the data. IR, HC, PJ and RAP wrote the paper.

## Material and methods

The morphometric analyses were done in Matlab using the TREES toolbox package (Cuntz et al., 2010). The statistical analsyis was done in R and the machine learning classification methods were implemented in Python. All data and code are made available at Zenodo upon publication but are available at the following Dropbox link for the reviewers:

https://www.dropbox.com/scl/fi/1f5xqwy6iknn9hp85v5xy/Raslain-et-al-CA2-radial-gradient.zip?rlkey=z88eiw3b8fyxrjzp1qni9x58r&st=q62l3yoh&dl=0

### Neuron reconstructions

128 CA2 PNs and 14 CA1 were recorded in whole-cell configuration in 400 *µ*m-thick transverse mouse hippocampal slices with biocytin in the intracellular pipette solution. Slices were fixed with 4% paraformaldehyde in phosphate buffered saline (PBS) overnight, washed with 0.3 M glycine in PBS for 20 minutes, then washed three times with PBS for 1 hour. Slices were permeabilized with 0.02% Triton in PBS and the biocytin was labeled with Alexa-598 conjugated strepavidin and neurons were labeled with 405-conjugated neurotrace. Confocal images were acquired as previously described (Robert et al., 2020; Chen et al., 2020; Robert et al., 2021). To obtain the 3D morphological data of CA1 and CA2 PNs, stacks of confocal images were analyzed with Neurolucida 360 (MicroBrightField). The reconstructed 3D morphological data were stored in an ASCII format and imported into MATLAB for further analysis.

### Importing the soma morphology

In Neurolucida 360, the soma is defined as a set of contours, and each contour is composed of points with different *X* and *Y* coordinates but the same *Z* coordinate. To align the somata with the rest of the neuronal tree, the orientation of these contours needed to be adjusted. This was achieved by finding the main axis of the somata points projected onto a 2D plane (without *Z* coordinates). The main axis was determined by identifying the two points that were farthest apart. The segment representing the main axis was then divided into regular intervals, and each set of points within these intervals was assigned a class. The basis was changed from the canonical basis to a new one, with the previously mentioned main axis becoming the Z axis in the 3D plane. The Z axis was then removed from each set of points, and three different methods were attempted to determine the largest radius of each contour: fitting an ellipse, fitting a circle, and obtaining the ferret diameter. The center of the selected shape for each contour was defined as the point in the somata tree structure, with the diameter dimension set to twice its corresponding radius. The root of the tree was determined as the point with the largest diameter. For somata volume determination, we used the 3D coordinates of the initial Neurolucida set of contours and applied the MATLAB function *convhull*. This approach provides an indirect assessment of the soma volume.

### Preprocessing

After importing the complete data set, several treatments were performed to enable a comparison between neurons. The neuronal trees obtained initially had varying distances between points, which could introduce bias in the analysis. To address this issue and ensure equal segment lengths for comparing specific metrics such as root angle distribution or cable length, a process known as resampling was employed. Resampling involved adjusting the distances between points to create standardized segment lengths, enabling more accurate and fair comparisons. First, the neuronal tree was rotated by aligning the main axis with the Y axis. Principal Component Analysis (PCA) was used for this purpose, and the first principal component (PC1) represented the main axis. Second, the neuronal tree was centered by subtracting the 3D coordinates of the root from all the coordinates of the tree. Finally, the tree was resampled into segments of 0.5 µm, using the *resample tree* function (Cuntz et al., 2010).

### Classifying neurons

To represent the boundaries of the layers, points connected by edges were used. In order to classify neurons, a gradient aligned with the Y-axis was drawn between the two boundaries that delineated the sp layer. This gradient ranged from 0 (indicating the so/sp boundary) to 1 (representing the sp/sr or sp/sl boundary). Neurons were classified based on the position of their root along this gradient called the soma location. Neurons with root falling between 0 and 0.5 were classified as deep neurons, while those falling between 0.5 and 1 were classified as superficial neurons.

### Defining pyramidal cell compartments

To define the basal and apical dendritic compartments, Neurolucida 360 provides the option to label them. However, for labeling the tuft, trunk, and oblique parts of the apical dendrite, we developed our own more objective algorithm. For CA2 PNs, our approach involved initially classifying the termination points of the apical dendrite into two clusters using the k-means algorithm from MATLAB (2022b). We obtained a distant cluster (from the root) representing the termination points of apical dendrites reaching the slm, and a closer cluster representing the termination points of apical dendrites ending in sr or sl. For each point in the distant cluster, we determined its direct path to the root using the *dA* matrix. We then identified the shared points among these paths and selected the first (from the terminations to the root) as the primary branch point. The apical dendrite reaching the slm from the primary branch point was defined as the tuft, while the part of the apical dendrite starting from the soma and ending at the primary branch point was defined as the trunk. The remaining portion of the apical dendrite constituted the oblique dendrites. In the case of CA1 PNs, we followed a similar procedure. However, we encountered challenges in classifying the termination points into two clusters using the *k-means* function as we did for CA2 PNs. Therefore, we adopted an alternative approach. By utilizing the *polyshape* function (MATLAB, 2022b) and the coordinates of the two boundaries delineating the slm layer, we constructed a polygon. Subsequently, by employing the *isInside* function (MATLAB, 2022b), we were able to classify the termination points of the neuronal tree. We then applied the same methodology as for CA2 PNs.

### Morphometrics

#### Sholl analysis

The Sholl intersection profile was generated using the *sholl_tree* function (Cuntz et al., 2010). To construct the profile, we determined the maximum Euclidean distance of the apical or basal dendrites and divided it into a vector of 25 equidistant values, starting from 0. These values were used as the radii for the Sholl Analysis. As a result, we obtained 25 data points representing the number of intersections at each radius. To analyze and compare the profiles, we used the *trapz* function (MATLAB, 2022b), which approximates the integral under the curve of the number of intersections as a function of the diameter using the trapezoidal method. This computation yielded 25 approximate integrals for both the apical and basal dendrites of both PN populations. To facilitate comparison, we scaled all 25 approximate integrals separately for apical and basal dendrites of both PN populations. The scaling was performed by normalizing the values between 0 and 1.

#### Root angle

The root angles were computed following the TREES toolbox *rootangle_tree* function, originally introduced by Bird and Cuntz (2019). To ensure uniformity, we resampled the dendritic arbor as mentioned in the preprocessing paragraph. Subsequently, the angle between the vector from the child to the parent and the vector from the child directly to the root was calculated. To determine the probability density function for each class of neurons, we applied the *pdf* function (MATLAB, 2022b). To determine the concentration parameters *κ*, we used the *vonMises tree* function also originally introduced by Bird and Cuntz (2019).

#### Branching order

The branching order of each point along the dendritic tree was determined using the *BO_tree* function (Cuntz et al., 2010). We then analyzed the data by counting the occurrences of each branch order value, resulting in a distribution of branching orders. To facilitate the comparison between conditions, we scaled the distributions individually for apical and basal dendrites. The scaling process involved normalizing the values within each distribution to a range between 0 and 1.

#### Oblique insertions

To compute the oblique insertions, we used the previously defined oblique compartment (**Methods** “Defining pyramidal cell compartments”) and looked for shared points with trunk or tuft-like compartments.

#### Cable length by section

To calculate the cable length of the dendrites, we used the *len_tree* function (Cuntz et al., 2010) on our resampled neuronal tree. This function provided us with the Euclidean distance between each child and parent point, allowing us to approximate the total length of the dendritic arbor by summing these distances. For each part of the apical dendrite, we calculated the proportional cable length (*CL*),

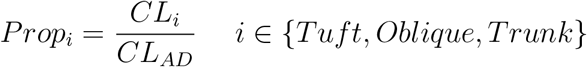

For the proportional apical dendrite cable length *CL*_*AD*_, we calculated the apical dendrite’s height defined as the distance from the root to the projection on the Y axis of the most distant termination point,

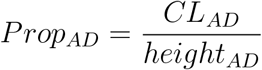

For the proportional basal dendrite cable length *CL*_*BD*_, we calculated the basal dendrite’s cable length and divided it by the total cable length of the tree. The Mean proportional *CL*, is the proportional *CL* of each section divided by the number of terminations points belonging to the section,

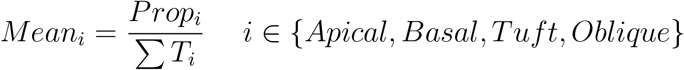

### Morphological classification using machine learning

To explore the potential classification or clustering of the CA2 PNs population into two subpopulations (dCA2 and sCA2), we employed machine learning techniques akin to those by Vasques et al. (2016). First, we computed various morphological measures for each neuron, encompassing concentration parameters of root angle distribution (apical, basal, tuft, oblique), maximum branching order (apical and basal), number of bifurcations (apical and basal), cable length (apical, basal, tuft, oblique, and trunk), number of terminations (tuft and oblique), proportional CL, mean proportional cable length by section (apical, basal, tuft, and oblique), and soma volume.

Next, for supervised learning, we selected algorithms based on their reported accuracy scores in prior research (Vasques et al., 2016), including k-nearest neighbors, linear discriminant analysis, support vector machine with a linear kernel, stochastic gradient descent, classification and regression tree, random forest, extremely randomized trees, and a multilayer perceptron. Each feature has been rescaled to a range between 0 and 1 using min-max normalization. This ensured that all features contributed equally to the model training. Each algorithm was evaluated using k-fold cross-validation with shuffling and a fixed random seed for reproducibility, to increase robustness, and to evade data overfitting. For each cross-validation, the following metrics were computed: accuracy, precision, recall, and F1-score. The mean and standard deviation of these metrics across all cross-validations were computed. CA2 data were shuffled and split into subsections of the same size as the CA1 data. This ensured balanced comparisons when applying the algorithms.

For unsupervised learning, we used algorithms by Vasques et al. (2016), including k-means, mini batch k-means, k-means on PCA-reduced data, Ward’s hierarchical clustering, mean shift, and affinity propagation. Before clustering, features were normalized using either L2 normalization (scaling to [−1,1]) or Z-score standardization (*mean* = 0, *std* = 1). To enhance model performance, we applied feature selection via recursive feature elimination with cross-validation (RFECV, optimized for accuracy) or random forest feature importance (threshold-based selection). Each algorithm was evaluated using homogeneity, completeness, V-measure (Rosenberg and Hirschberg, 2007), adjusted Rand index, and silhouette score. Hyperparameter tuning was performed for RFECV, maximizing cross-validation accuracy, and for mean shift, k-means (PCA-reduced), as well as for affinity propagation, maximizing V-measure.

All algorithms were implemented using Python 3.11.5, leveraging the Scikit-learn open-source Python library.

### Statistics

Analyses were conducted on a dataset comprising 14 CA1 PNs and 128 CA2 PNs. We conducted the Shapiro-Wilk test to assess normality and the Fisher-Snedecor test to examine homoscedasticity, allowing us to determine whether parametric or non-parametric statistical analyses were suitable. To evaluate the statistical significance of variables, we employed the unpaired Student’s t-test and Mann–Whitney-Wilcoxon U-test as appropriate for our analysis. Results are reported as mean ± SEM. Additionally, to investigate the correlation between soma location and morphological tree measures, we calculated correlation coefficients using the Pearson or Spearman correlation test and computed the moving average. All statistical tests were conducted using the R environment (R Core Team 2022): *shapiro*.*test, var*.*test, t*.*test, wilcox*.*test, cor*.*test*.

